# Electrical charge-mediated transformation for efficient plant genome editing

**DOI:** 10.1101/2024.11.19.624257

**Authors:** Sruthy Maria Augustine, Anoop Vadakan Cherian, Paridhi Paridhi, Md Mamunur Rashid, Samson Ugwuanyi, Babette Knoblauch, Stavros Tzigos, Soni Savai Pullamsetti, Rod Snowdon

## Abstract

Genome editing technologies possess significant potential to enhance plant breeding; however, the delivery of editing constructs poses challenges in numerous crop species due to their resistance to transformation or tissue culture. This study introduces an innovative approach for the delivery of ribonucleoprotein (RNP) complexes and plasmid vectors into intact, regenerable faba bean plant tissues. This is accomplished by applying an electric current that makes plant cell walls and membranes permeable, thereby allowing the entry of macromolecular constructs into the cell and nucleus. This study assessed the efficacy of electric pulse-mediated transfection in faba bean by generating stable GFP-expressing faba bean plants. Furthermore, we incorporated it into faba bean leaf tissue and demonstrated its application in both embryos and leaf tissues. We demonstrate DNA-free genome editing by targeting the endogenous phytoene desaturase gene (PDS), achieving a mutation success rate of 50%. This method is efficient and economical, necessitating limited technical training. It is applicable to both leaf and embryo tissues, thereby enhancing its utility for crop improvement. This technique shows potential for the development of new crop varieties that can more effectively address global climate challenges.

---

Genome editing technologies have great potential to accelerate plant breeding, but delivery of editing constructs is difficult in many crop species because they are recalcitrant to transformation or tissue culture. Here, we present a transformative method for delivering ribonucleoprotein (RNP) complexes and plasmid vectors into intact, regenerable plant tissues. This is achieved by applying an electric current to make plant cell walls and membranes permeable, facilitating the entry of macromolecular constructs into the cell and nucleus.

We demonstrate the utility of the method for genome editing in faba bean (*Vicia faba* L.), one of the earliest domesticated crops and a significant cool-season legume in global agriculture, valued for its nitrogen-fixing capabilities and role as a primary protein source in many countries (Jithesh et al., 2024; Jayakodi et al., 2023). A major limitation in faba bean research to date has been the lack of a reliable transformation or genome-editing method. In preparation, the excised seed embryo is perforated 5-8 times with a 26-gauge needle, followed by application of a droplet of liquid containing the desired macromolecular construct, such as an RNP complex or plasmid, onto the perforated surface. Two 26-gauge needles are then inserted into the plant tissue, and transfection is achieved by applying an electric current via a 24-28 V battery (Figure 1A).

**Figure 1:**
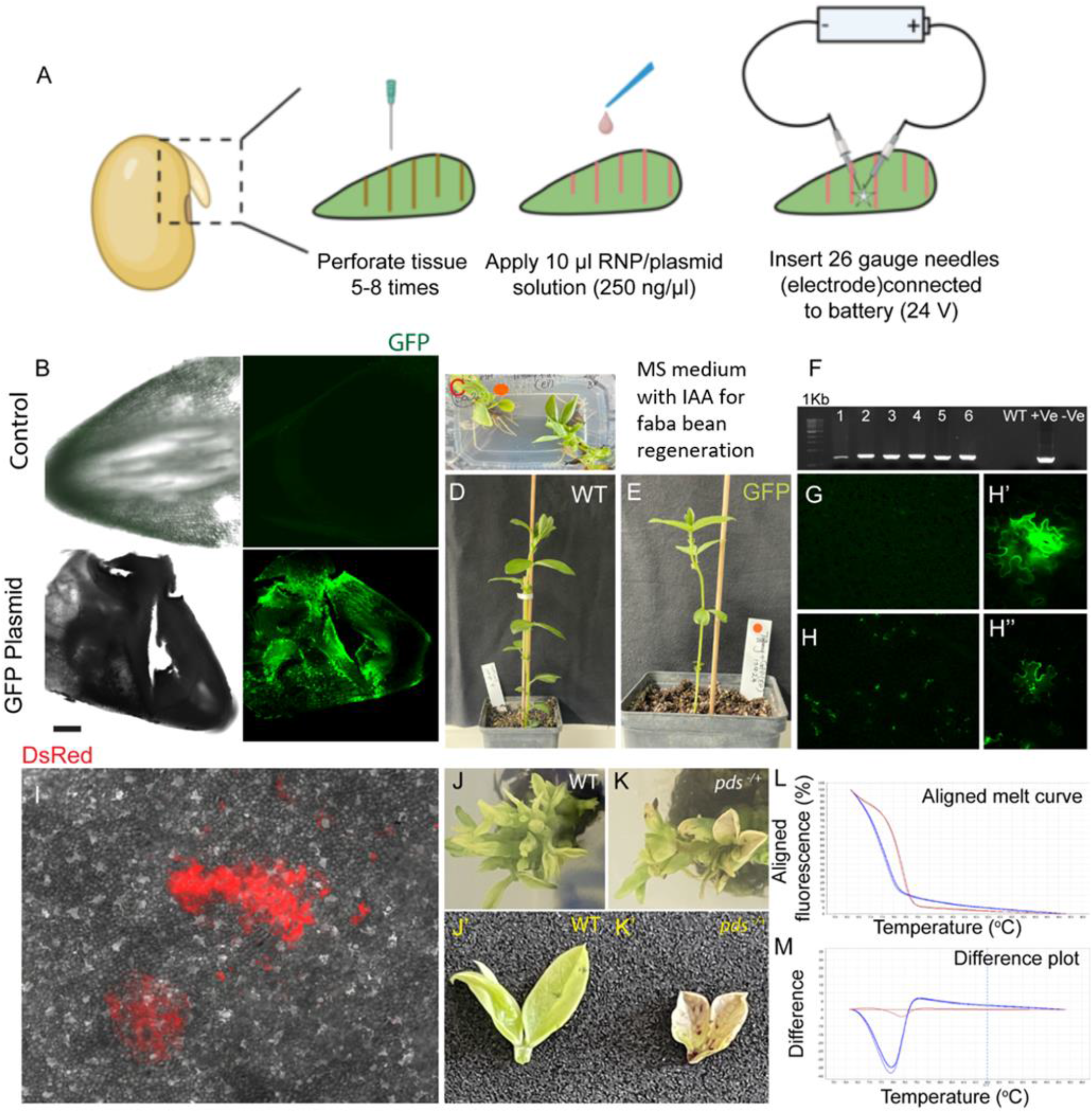
Electric charge mediated genome editing in faba bean. **(A)** Electric pulse-mediated transfection of an intact plant embryo via battery-connected needle electrodes. **(B)** Verification of electric pulse-mediated transformation of *Vicia faba* embryos with the pLH-6000-GFP construct by evaluation of GFP expression in the wild type control (above) and transfected embryo (below) two days after transfection. **(C)** MS medium with IAA for faba bean regeneration. **(D)** Regenerated wild type control. **(E)** Regenerated GFP overexpressed faba bean. **(F)** PCR confirmation of the hpt gene in the GFP overexpressed faba bean plants. (G) GFP analysis in wild type control. **(H)** GFP expression in leaf tissue of the GFP overexpressed plant. **(H’, H”)** Zoomed regions from Figure H. **(I)** DsRed fluorescence detection on the faba bean leaf tissue after electric charge mediated transfection. **(J, J’)** Wild type faba bean plants. **(K, K’)** Chimeric albino mutant plants resulting in mutation of the *pds* gene. High-resolution melt analysis of mutant plants generated through gene editing by electric charge-mediated transfection resulting in mutation of the pds gene **(L)** Aligned melt curve **(M)** Difference plot.

To evaluate the effectiveness of electric pulse-mediated transfection in faba bean, we introduced the green fluorescent protein (GFP)-expressing transgene construct pLH-6000-GFP (Figure S1; Imani et al., 2011) into faba bean embryos using this technique. Embryos were extracted from mature seeds that had been soaked in sterile MilliQ water for around 16 hours. Viable embryos were retrieved and transferred to a standard 100×15 mm petri dish. One needle was inserted 2-3 mm into the embryo, while the second was positioned adjacent to the first, creating an electric charge to enable plasmid transfection into the cells. GFP presence was confirmed two days post-electric pulse using confocal microscopy (Figure 1B), with fluorescence detected using a 488 nm laser for excitation and an emission peak near 508 nm. Post-transfection, embryos expressing GFP were cultured on MS medium (4.4 g/L MS salts with vitamins, 20 g/L sucrose, 7 g/L agar, pH 5.8) supplemented with 1.5 mg/L IAA and maintained at 20-23°C. The regenerated tissue was subcultured every 10-12 days until shoot emergence (Figure 1C). The plants were then transferred to greenhouse conditions alongside wild-type controls (Figure 1D and 1E). Genomic DNA was extracted at the 4-6 leaf stage following Doyle and Doyle (1990), and the presence of the hygromycin (hpt) marker gene was confirmed via PCR (Figure 1F). The transgenic status was further validated by observing GFP expression in leaves with confocal microscopy (Figure 1G, H, H’, H”).

In addition to embryos, we assessed the efficacy of this method for introducing constructs and RNPs into leaf tissues using 3-4-month-old faba bean leaves. The technique successfully incorporated a red fluorescent protein (DsRed) R2G mutant from pGJ1425 (MPI, Cologne, Germany) (Sack et al., 2015) into leaf tissue, indicating that this method is adaptable to both embryos and leaf tissues as explants (Figure 1I).

To demonstrate DNA-free genome editing, we targeted the endogenous phytoene desaturase gene (PDS), where mutations yield a visually identifiable albino phenotype, providing an efficient measure of mutation success. The gene structure is shown in Figure S2, with the gene identified as *Vfaba*.*Tiffany*.*R1*.*2g090080* and the corresponding transcript *Vfaba*.*Tiffany*.*R1*.*2g090080*.*1* (Tiffany sequence available at https://projects.au.dk/fabagenome/genomics-data, Jayakodi et al., 2023). Target regions within the *PDS* gene were amplified from genomic DNA using Q5 high-fidelity DNA polymerase, purified with the QIAquick gel extraction kit (QIAGEN, Hilden, Germany), and sequenced using Sanger sequencing before crRNA design, as specified in Supplementary Table 1. crRNA sequences were generated using the online tool “CRISPRdirect” (https://crispr.dbcls.jp/). The crRNA targeting the *PDS* gene was 5’-GAACCATGGTTCTCGTTTGA-3’.

RNP complexes were prepared with crRNA, tracrRNA, and Cas9 protein (synthesized by Integrated DNA Technologies, Inc., Iowa, USA). For all experiments, chemically modified crRNA-XT was used to enhance stability and performance. Equimolar crRNA and tracrRNA were mixed at a final concentration of 100 μM and heated at 95°C for 5 minutes. Next, 120 pmol of gRNA mix, 104 pmol of Cas9 protein, and 2.1 μL of PBS were combined for a total RNP volume of 5 μL and incubated at room temperature for 20 minutes. The RNP complex was then immediately administered to freshly isolated embryos, eliminating the need for labor-intensive protoplast or zygote preparations. Following RNP delivery, embryos were cultured on MS medium at 20-23°C without selection reagents. Chimeric albino mutants from *PDS* gene knockouts were identified visually (Figure 1K, K’, S3), with wild-type controls shown in Figure J and J’. Following regeneration, high-resolution melting analysis (HRMA), a PCR-based method, was used to detect chimeric and heterozygous mutants (Denbow et al., 2018; Li et al., 2018). In the analysis of 22 plants from the *PDS* gene editing experiment, 11 exhibited mutations at the T0 stage. Figure 1L and M show the melting curves of *PDS* mutants compared to wild-type controls, with additional HRMA curve images provided in Figure S4. All amplicons measured 100-150 bp in length, with conditions and primers detailed in Supplementary Table 2.

Our electric pulse-mediated transformation achieved a 50% mutation efficiency in the *PDS* gene within approximately 7-8 months. These findings suggest that the electric charge-mediated transformation method is effective for genome editing in recalcitrant species like faba bean. Direct transformation or mutation of intact embyros can overcome difficulties in large-seeded plant species which are recalcitrant to tissue culture regeneration from alternative explants. This straightforward, cost-effective method, which requires minimal technical training, is applicable to both leaf and embryo tissues, making it broadly useful for crop improvement. This technique holds promise for developing new crop varieties that can better respond to global climate challenges.

## Author contributions

SMA and AVC designed the study, conducted data analyses, and wrote the manuscript. PP, MMR, BK, and ST performed and provided technical assistance in the work; SU provided the bioinformatics support in the study; SSP provided the confocal microscope facility; and RS conceptualised and critically reviewed the manuscript. All authors have reviewed and approved to the final manuscript.

## Acknowledgement

The authors thank Norddeutsche Pflanzenzucht Hans-Georg Lembke KG (NPZ, Hohenlieth, Germany) for providing the faba bean seeds used in the study.

## Supplementary Information

### Supplementary figures

**Figure S1:**
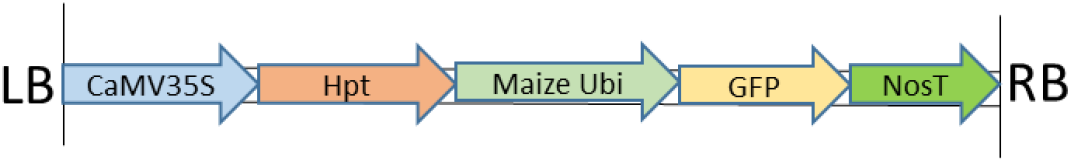
pLH-6000-GFP construct map

**Figure S2:**
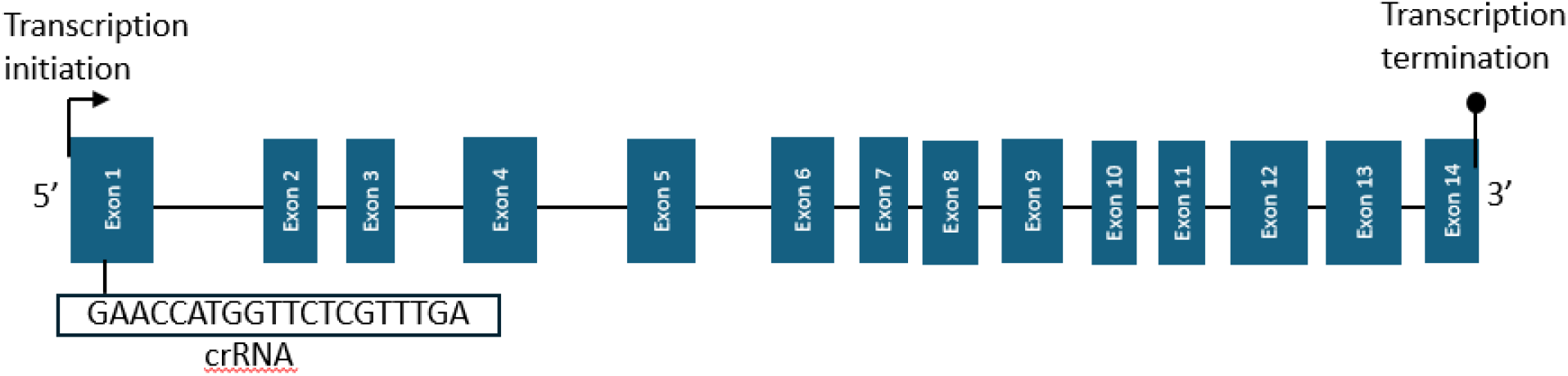
Schematic representation of the structure of faba bean PDS gene, showing the exons and introns. The blue boxes represent the exons, while the dark lines represent the introns. The width of the boxes corresponds to the length of the exons, and the length of the lines between the boxes corresponds to the length of the introns. Position and sequence of CRISPR RNA (crRNA) are shown above.

**Figure S3:**
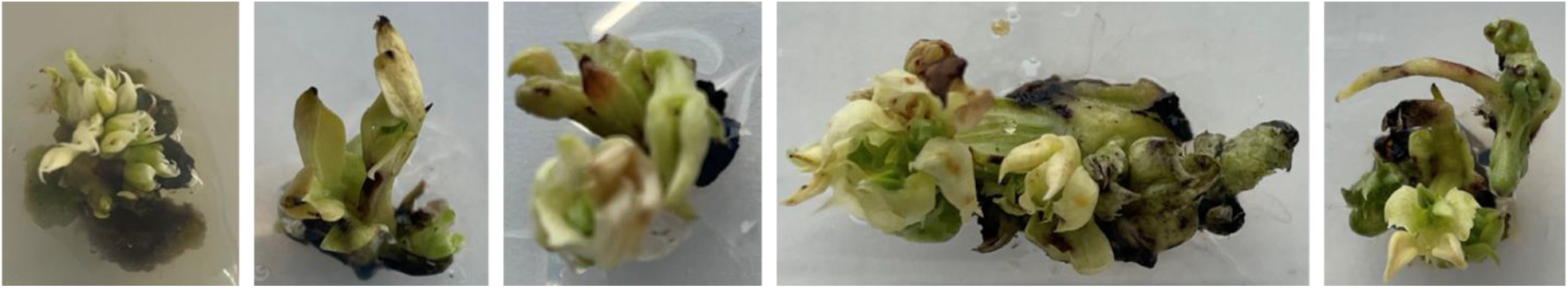
Albino mutant plants after gene editing by electric pulse-mediated transfection resulting in mutation of the pds gene.

**Figure S4:**
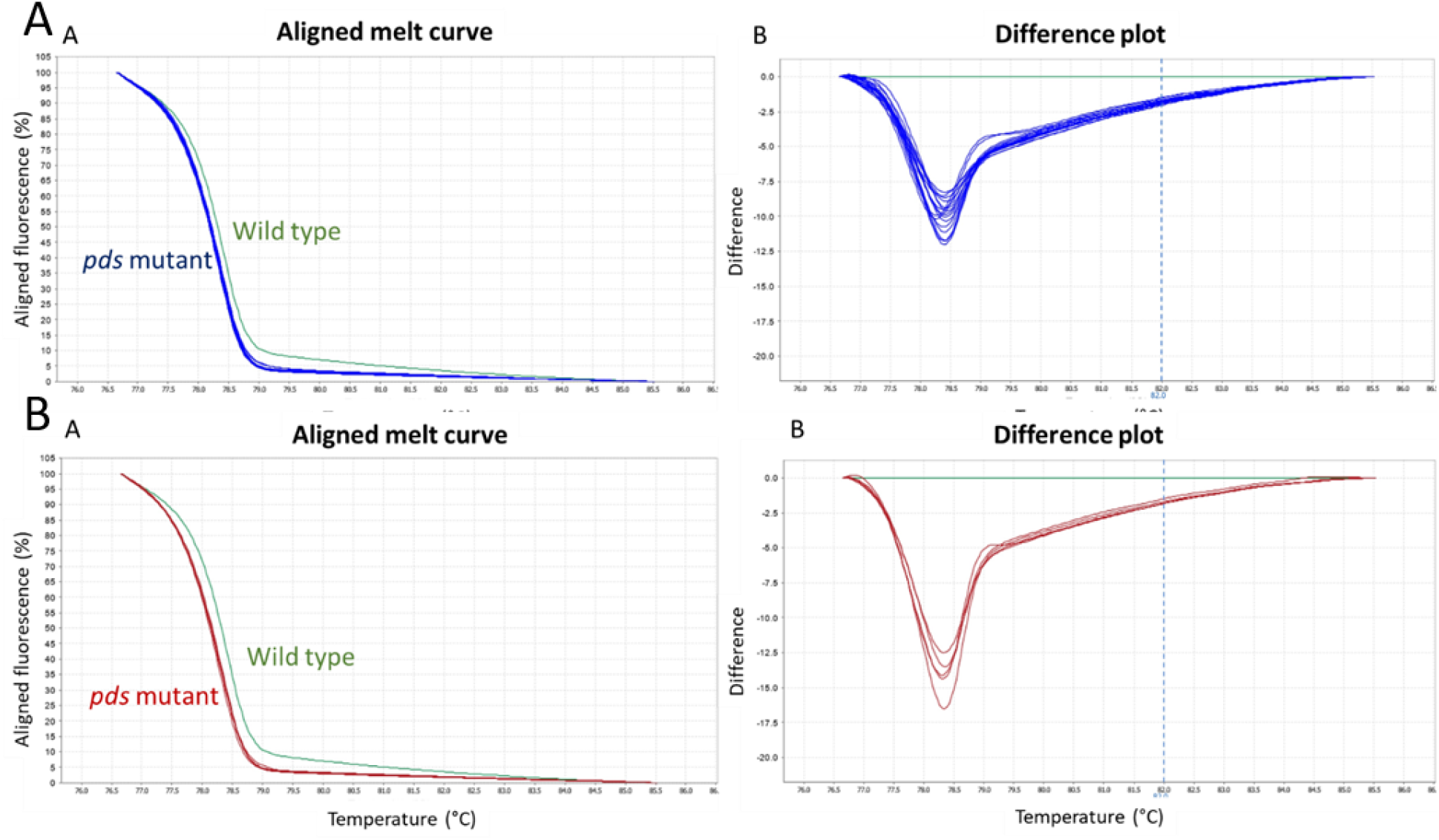
High-resolution melt analysis of mutant plants generated through gene editing by electric pulse-mediated transfection resulting in mutation of the pds gene. A) A) Aligned melt curve showing the difference in pds mutant and wild type. B) Difference plot.

### Supplementary tables

**Supplementary Table 1:**
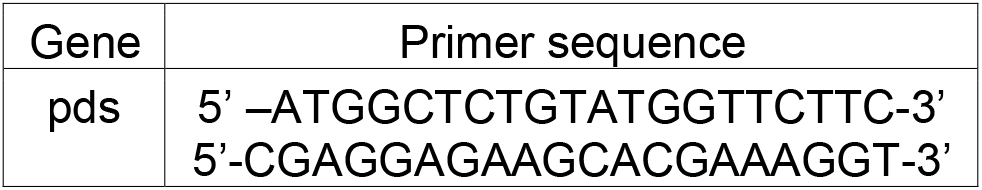
pds gene primer sequence.

**Supplementary Table 2:**
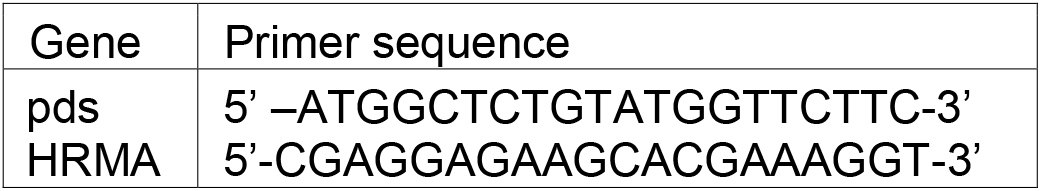
pds gene HRMA amplification protocol. For the PDS gene, the following conditions were used: PCR stage, 95°C for 10 min, followed by 40 cycles of 95°C for 15 s, 52°C for 1 min, and 72°C for 1 min; melt curve stage, 95°C for 15 s (1.6°C/s), 60°C for 1 min (1.6°C/s) and 95°C for 15 s (0.1°C/s).

